# From SNPs to Pathways: A genome-wide benchmark of annotation discrepancies and their impact on protein- and pathway-level inference

**DOI:** 10.64898/2026.03.21.713397

**Authors:** Bryan Queme, Anushya Muruganujan, Dustin Ebert, Tremayne Mushayahama, W. James Gauderman, Huaiyu Mi

**Author notes:** **Corresponding Author:** Huaiyu Mi.

## Abstract

**Background:** Accurate single-nucleotide polymorphism (SNP) annotation is central to genomic research yet widely used tools and gene models often yield divergent results. Prior studies have shown such discrepancies in small datasets, but the extent of genome-wide variation and its impact on downstream pathway analysis remain unclear.

**Results:** We conducted a comprehensive comparison of three commonly used SNP annotation tools, ANNOVAR, SnpEff, and VEP, using both Ensembl and RefSeq gene models to evaluate more than 40 million SNPs from the Haplotype Reference Consortium. At the protein level, annotation output differed significantly across tools and gene models (p-adj < 0.001), with discrepancies present in both genic and intergenic regions. RefSeq produced broader annotation coverage, particularly for intergenic SNPs, while Ensembl showed greater internal consistency. SnpEff provided the most complete coverage overall, whereas no single tool or model configuration achieved full annotation recovery of the union reference. Integration across tools and models maximized coverage and reduced annotation loss. In a case study of 204 colorectal cancer–associated SNPs from the FIGI GWAS, pathway enrichment results varied depending on annotation strategy. The fully integrated approach identified all four significant pathways, whereas several single-tool or single-model strategies missed one or more.

**Conclusion:** SNP annotation outcomes are influenced by both the tool and gene model used, and relying on a single approach may result in incomplete coverage. A multi-tool, multi-model strategy provides the most comprehensive annotation and preserves enriched pathways, supporting more robust and reproducible genomic interpretation.

## BACKGROUND

Single-nucleotide polymorphisms (SNPs), the most prevalent form of genetic variation in the human genome, play a central role in understanding disease susceptibility^1^, gene regulation^2^, and phenotypic diversity^3^. Interpreting SNPs accurately depends on gene annotation tools that map each variant to one or more genes and predict how those variants might affect gene or protein function, which is an essential step in studies of complex traits^4^, pharmacogenomics^5^, and disease etiology^6,7^.

A common application of SNP annotation is biological pathway analysis^8^, which allows researchers to assess whether coding genes (or their protein products), linked to phenotype-associated SNPs, such as those identified in case-control or genome-wide association studies, are disproportionately represented in specific biological pathways^9^. This approach involves mapping SNPs to protein-coding genes, which are then compared against curated pathway databases like Reactome^10^ and PANTHER^11,12^ using pathway enrichment methods^8,13,14^. Because these analyses depend on the accuracy of SNP-to-protein mapping, pathway results can be strongly influenced by the annotation tools^15,16^ or gene models^17,18^.

A wide range of tools have been developed for SNP annotation, including ANNOVAR^19^, SnpEff^20^, and the Variant Effect Predictor (VEP)^21^. These tools rely on reference gene models, curated definitions of gene and transcript boundaries, such as Ensembl^22^ and RefSeq^23^, which differ in how genes and transcripts are defined and annotated. Ensembl includes more alternative transcripts and regulatory features, while RefSeq emphasizes curated gene models^17^. These structural differences can yield divergent interpretations of the same SNPs. Previous studies have noted such discrepancies using focused datasets or transcriptomic contexts, but to our knowledge, none have quantified this variation across the entire genome using standardized inputs and outputs^15–18^.

To address this gap, we conducted a comprehensive, genome-wide assessment of annotation consistency across ANNOVAR, SnpEff, and VEP, using both Ensembl and RefSeq gene models. Leveraging over 40 million SNPs from the Haplotype Reference Consortium (HRC)^24^, we compared annotations across tools and gene models in both genic regions, within or near genes, and intergenic regions, between genes. Although intergenic SNPs do not alter coding sequence, they frequently map to regulatory regions or nearest genes and are routinely included in post-GWAS functional analysis^25,26^.

We used two complementary approaches to evaluate agreement at the SNP level: (1) **a qualitative analysis of annotation overlap patterns for each SNP**, such as whether annotations were tool-specific or shared across tools, and (2) **a quantitative analysis measuring the proportion of protein-coding annotations captured by each tool per SNP**, relative to a unified reference set.

Furthermore, to evaluate these downstream effects, we applied our annotation framework to a pathway enrichment case study using 204 SNPs from the Functionally Informed Gene–Environment Interaction (FIGI) colorectal cancer GWAS^27,28^. Our findings indicate that the choice of annotation tool and gene model substantially influences both annotation coverage and pathway enrichment results. By systeatically quantifying these discrepancies using a consistent SNP input set and protein-level outputs, this study provides the first genome-wide assessment of tool- and gene model-driven variation in SNP-to-protein annotation, offering practical guidance for improving reproducibility and interpretability in SNP-based genomic research.

## RESULTS

### Annotation Overview

While annotating the 40,290,938 unique SNPs from the HRC dataset, we observed substantial differences between Ensembl and RefSeq in the number of SNPs successfully mapped to UniProt IDs when aggregating annotations across all three tools within each gene model (Figure 1, Table 1).

**Figure 1.**
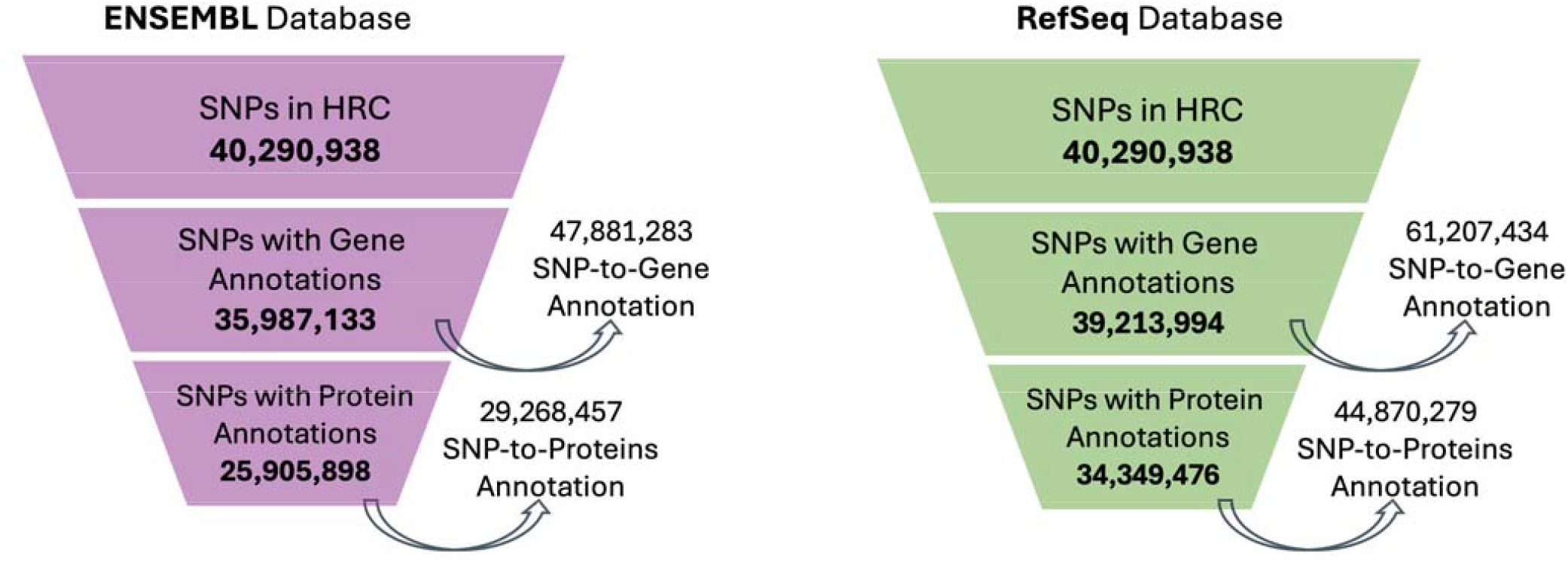
Stepwise reduction of SNPs with gene and protein annotations across Ensembl and RefSeq. This funnel plot illustrates the annotation trajectory of 40,290,938 SNPs from the Haplotype Reference Consortium (HRC), first to gene annotations and then to UniProt protein identifiers. Each layer shows the number of SNPs with at least one successful annotation at that level. Side arrows reflect total mappings, accounting for cases where individual SNPs map to multiple genes or proteins. RefSeq annotated more SNPs at both gene and protein levels than Ensembl, highlighting systematic differences in annotation yield and gene model scope.

**Table 1.** Protein Annotation Coverage by Gene Model for Each Tool, Using Standardized Region Definitions. Percentage of UniProt IDs captured by each tool (ANNOVAR, SnpEff, VEP) when using RefSeq versus Ensembl gene models, with region definitions standardized to Ensembl. Values represent the proportion of the SNP-specific UniProt reference set annotated by each tool and gene model combination, stratified by region (whole genome, genic, intergenic). This table isolates the effect of gene model selection while holding the annotation tool constant and highlights how RefSeq generally yields higher coverage than Ensembl, except for VEP, where the trend is reversed. Because the ‘Combined’ column is the union reference for each tool across gene models under the standardized framework, it attains 100% recovery by definition; the informative quantities are the shortfalls of the single-gene-model configurations (RefSeq or Ensembl) relative to that union.

| Gene Models |  |  |  |
| --- | --- | --- | --- |
|  | RefSeq | Ensembl | Combined |
| Genic |  |  |  |
| ANNOVAR | 24,138,918 (95.0%) | 20,008,036 (78.7%) | 25,422,674 (100%) |
| SnEff | 25,811,247 (94.0%) | 21,451,709 (78.1%) | 27,461,035 (100%) |
| VEP | 18,775,184 (90.4%) | 20,318,587 (97.8%) | 20,777,302 (100%) |
| Intergenic |  |  |  |
| ANNOVAR | 17,456,051 (96.3%) | 7,394,112 (40.8%) | 18,119,063 (100%) |
| SnEff | 18,368,241 (96.0%) | 7,786,423 (40.7%) | 19,128,052 (100%) |
| VEP | 226,680 (50.2%) | 275,070 (60.9%) | 451,729 (100%) |
| Whole |  |  |  |
| ANNOVAR | 41,594,969 (95.5%) | 27,402,148 (62.9%) | 43,541,737 (100%) |
| SnEff | 44,179,488 (94.8%) | 29,238,132 (62.8%) | 46,589,087 (100%) |
| VEP | 19,001,864 (89.5%) | 20,593,657 (97.0%) | 21,229,031 (100%) |

Ensembl annotated 35,987,133 SNPs to at least one gene, collectively mapping 47,881,283 SNP-o-Gene annotations, reflecting instances where SNPs mapped to multiple genes. Of those, 25,905,898 SNPs were successfully linked to at least one UniProt ID, accounting for a total of 29,268,457 SNP-to-Protein annotations (Figure 1).

In contrast, RefSeq yielded gene annotations for 39,213,994 SNPs, corresponding to 61,207,434 SNP-to-Gene annotations. From these, 34,349,476 SNPs were mapped to at least one UniProt ID, generating 44,870,279 SNP-to-Protein annotations.

Figure 2 illustrates how using Ensembl and RefSeq native gene boundaries differs in the proportion of SNPs classified as genic versus intergenic based on ANNOVAR’s classification^19^. Out of the 40,290,938 SNPs in the HRC dataset, annotations using Ensembl gene boundaries classified 61.5% as genic and 38.5% as intergenic. Using RefSeq gene boundaries, the proportions were 51.2% genic and 48.8% intergenic. However, among SNPs that were successfully mapped to at least one protein, these proportions differed. For Ensembl, genic SNPs comprised 61.5% of all SNPs but 75.2% of SNPs that could be mapped to at least one protein, indicating preferential retention of genic variants during protein mapping. RefSeq, in contrast, maintained a nearly identical genic-to-intergenic ratio (53.0% genic, 47.0% intergenic) among its 34,349,476 protein-annotated SNPs compared to 51.2% genic and 48.8% intergenic.

**Figure 2.**
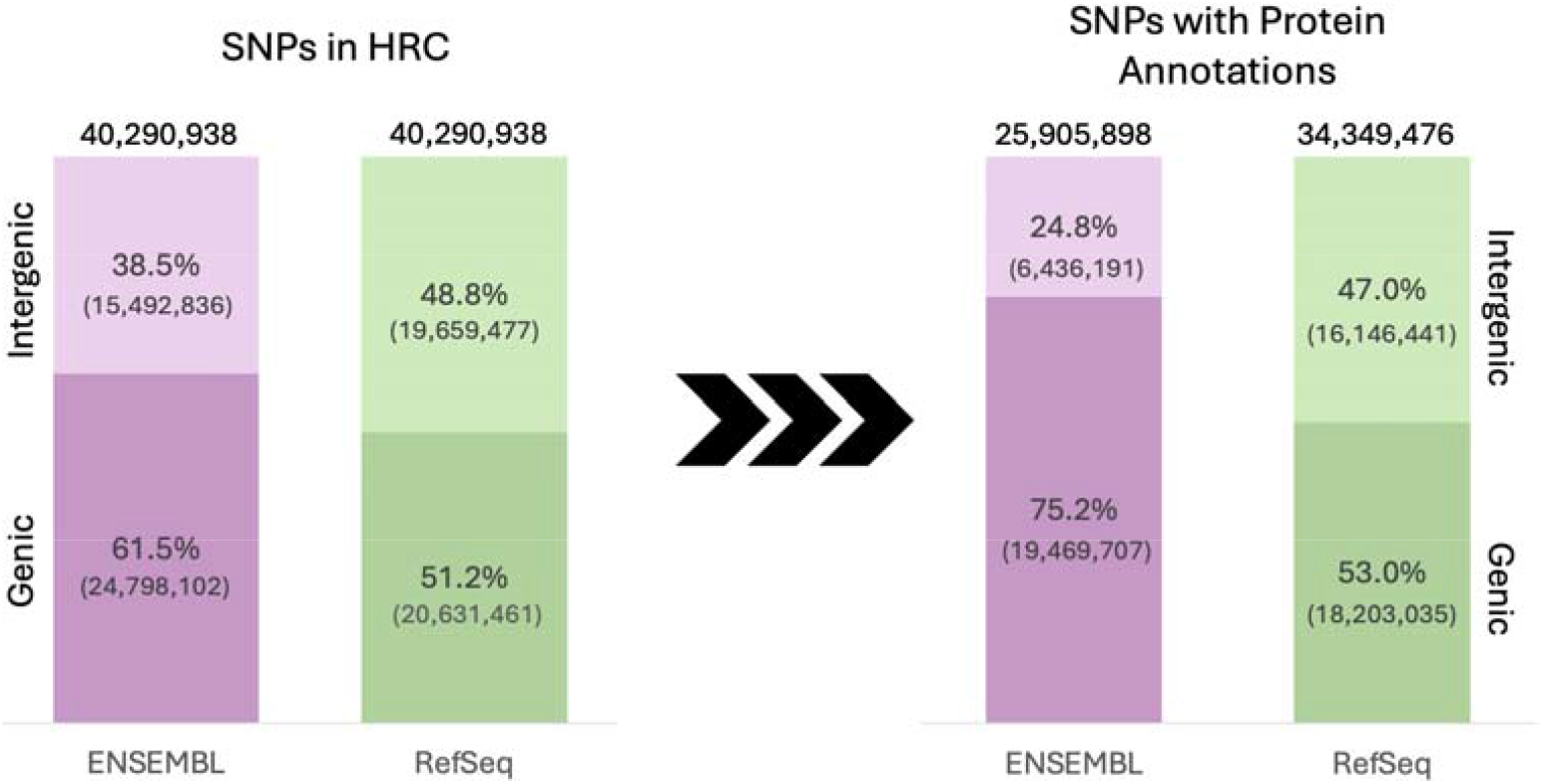
Gene model-level differences in genic vs. intergenic region assignment. Stacked bar plots show the proportion of SNPs annotated to genic and intergenic regions using Ensembl and RefSeq, based on ANNOVAR definitions. Left: All 40,290,938 SNPs from the HRC dataset. Right: Subset of SNPs with at least one UniProt protein annotation, 25,905,898 for Ensembl and 34,349,476 for RefSeq. Ensembl assigns a larger proportion of total SNPs to genic regions (61.5%) than RefSeq (51.2%). Among protein-annotated SNPs, this difference widens, 75.2% of Ensembl-mapped SNPs are genic versus 53.0% for RefSeq. These patterns suggest differences in annotation scope and prioritization between gene models, especially at the protein level.

These results indicate that RefSeq provides broader protein annotation coverage, capturing 32.6% more SNPs at the protein level than Ensembl (34,349,476 vs 25,905,898 SNPs, respectively) and preserves genomic region proportions more consistently during annotation. This contrast highlights how different gene models influence not only overall coverage but also the balance of genic and intergenic signals in downstream analyses.

These discrepancies reflect inherent structural and content differences between the two gene models, emphasizing the impact of model selection on downstream annotation depth. To distinguish tool-from gene model-driven effects, we next evaluated agreement patterns among tools and between gene models.

### Benchmarking Annotation Agreement

To evaluate the consistency with which annotation tools and gene models assign functional consequences to SNPs, we applied two complementary approaches at the SNP level (see the Annotation Ag eement Framework section). First, we used a **qualitative analysis** to assess whether tools agreed at all on which proteins were assigned to each SNP. This analysis focuses on overlap patterns, such as whether annotations were tool-specific or shared, and highlights structural differences between tools and gene models. Second, we used a **quantitative analysis** to assess how completely each tool recovered the full set of protein annotations for each SNP, using the proportion of UniProt IDs captured relative to a unified reference set. Together, these analyses provide both a high-level view of agreement patterns and a detailed measure of annotation completeness.

We first examined **qualitative agreement** between annotation tools. Using each gene model’s native region definitions from ANNOVAR, across the genome, we found that all three tools, ANNOVAR, SnpEff, and VEP, produced identical protein annotation sets for only 69.9% of SNPs using Ensembl and 47.3% using RefSeq (Additional File 2). When both gene models were combined, three-tool agreement reached 48.7%, indicating a modest increase compared to RefSeq alone but still substantially lower than the agreement observed using Ensembl.

The next most common pattern was agreement between ANNOVAR and SnpEff only, without any annotations from VEP. This category accounted for 23.0% of Ensembl SNPs, 41.7% of RefSeq SNPs, and 39.2% of SNPs in the combined gene model configuration. Other patterns, including subset relationships or agreement between VEP and a single other tool, were relatively rare (Additional File 2).

To determine whether tool agreement varied between genomic contexts, we repeated the analysis separately for genic and intergenic regions. In genic regions, three-tool agreement was high across all configurations: 92.7% of Ensembl SNPs, 88.8% of RefSeq SNPs, and 75.6% when annotations from both gene models were combined. In contrast, intergenic agreement was far lower: only 0.8% for Ensembl, 0.5% for RefSeq, and 1.2% in the combined gene model set. The dominant pattern in intergenic regions was agreement between ANNOVAR and SnpEff only, which accounted for 92.4% of Ensembl SNPs, 88.7% of RefSeq SNPs, and 87.2% of SNPs in the combined model configuration.

Finally, we assessed annotation agreement between gene models. For each SNP, we took the union of annotations across tools within Ensembl and RefSeq separately. We compared the resulting protein annotations using ANNOVAR-derived region definitions based on the Ensembl gene model (Additional File 3). Genome-wide, only 56.2% of SNPs had identical protein annotations between gene models. RefSeq-only annotations accounted for 26.4%, and cases where RefSeq was a strict superset of Ensembl made up 11.7%. In genic regions, agreement rose to 73.8%, while “RefSeq-only” and “RefSeq with Ensembl Subset” categories accounted for 13.3% and 6.9%, respectively. Discrepancies were especially pronounced in intergenic regions, where exact matches occurred in just 25.2% of SNPs and RefSeq-only annotations accounted for 49.5% (Additional File 3).

This qualitative analysis provides a detailed SNP-level view of annotation agreement, capturing whether different tools and gene models assigned the same or overlapping proteins to each variant. By examining each SNP individually, we were able to identify nuanced patterns of agreement and tool-specific behavior that would be obscured in aggregate counts. Having established qualitative differences in how tools assign proteins to individual variants, we next quantified annotation completeness. This quantitative analysis aggregates per-SNP coverage at the chromosome level to support statistical comparisons across tools and gene models.

The **quantitative agreement analysis** focused on how each tool recovered the total set of protein annotations per SNP (Figure 3). We calculated, for each tool, the proportion of UniProt IDs it captured out of a unified reference set and aggregated these values at the chromosome level to allow statistical comparison. Similarly to the qualitative agreement analysis, we were interested in comparing within and across gene models.

**Figure 3.**
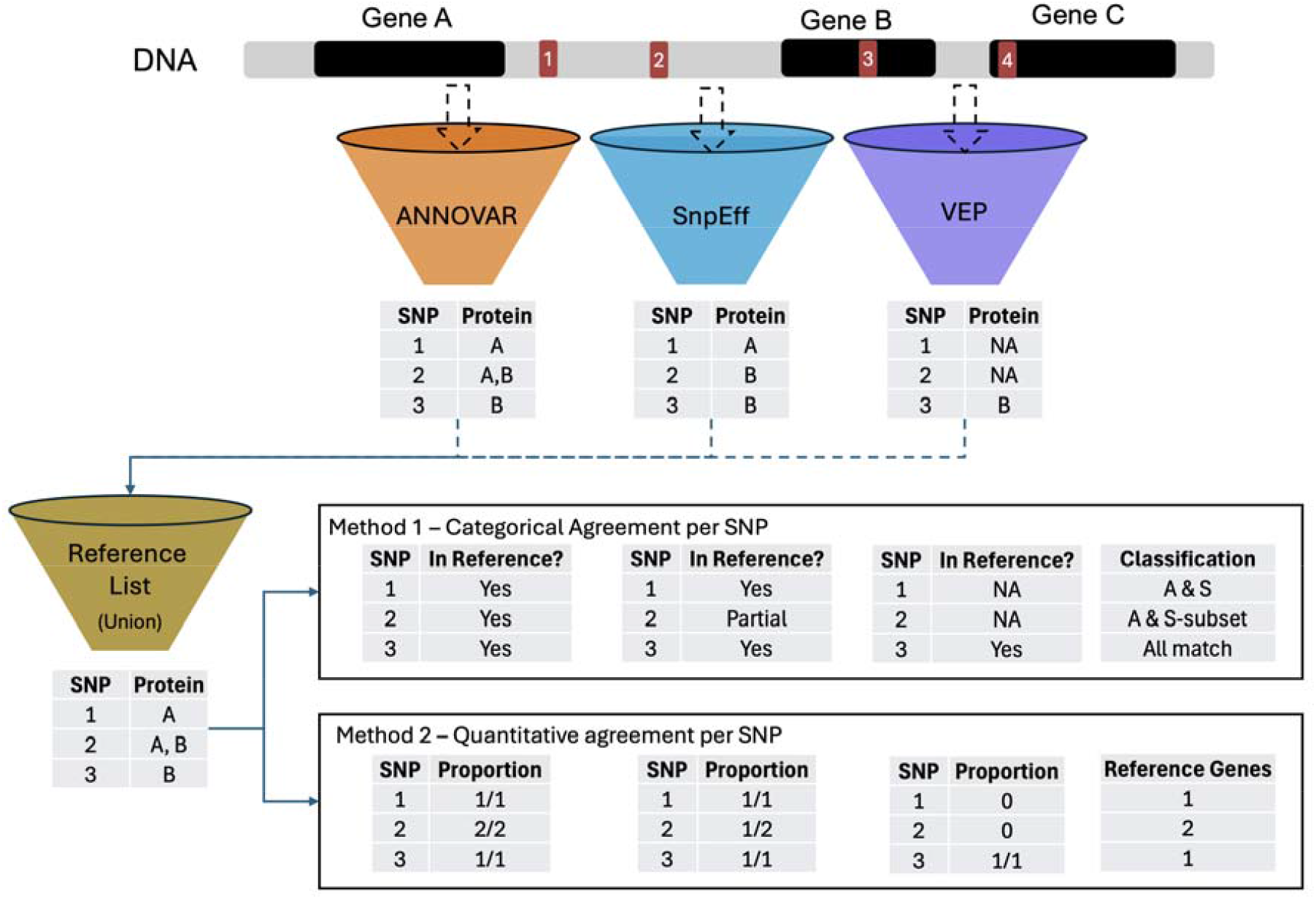
Overview of annotation agreement framework for protein-level SNP mapping. SNPs from the HRC dataset were annotated using three tools (ANNOVAR, SnpEff, and VEP), each returning one or more gene assignments per SNP. For each SNP, a reference protein set was defined as the union of protein annotations across tools. Categorical agreement was assessed by determining whether each tool’s annotations fully, partially, or failed to match the reference set; these patterns were used to assign each SNP to a descriptive agreement class (e.g., “ANNOVAR and SnpEff,” “ANNOVAR; SnpEff subset,” “Exact Match”). Qantitative agreement was recorded as the number of reference proteins captured by each tool (e.g., 2 out of 3), and these counts formed the basis for subsequent statistical analyses. A comparable framework was used to evaluate agreement across gene models (Ensembl and RefSeq), substituting tools with models in the reference logic.

Within RefSeq and Ensembl, all pairwise comparisons between tools were statistically significant after Bonferroni correction (p-adj < 0.001), with one exception (Fig. 4A-C; 5A, 5C): ANNOVAR and VEP did not significantly differ in RefSeq genic regions (p-adj > 0.05; Fig. 5B).

**Figure 4.**
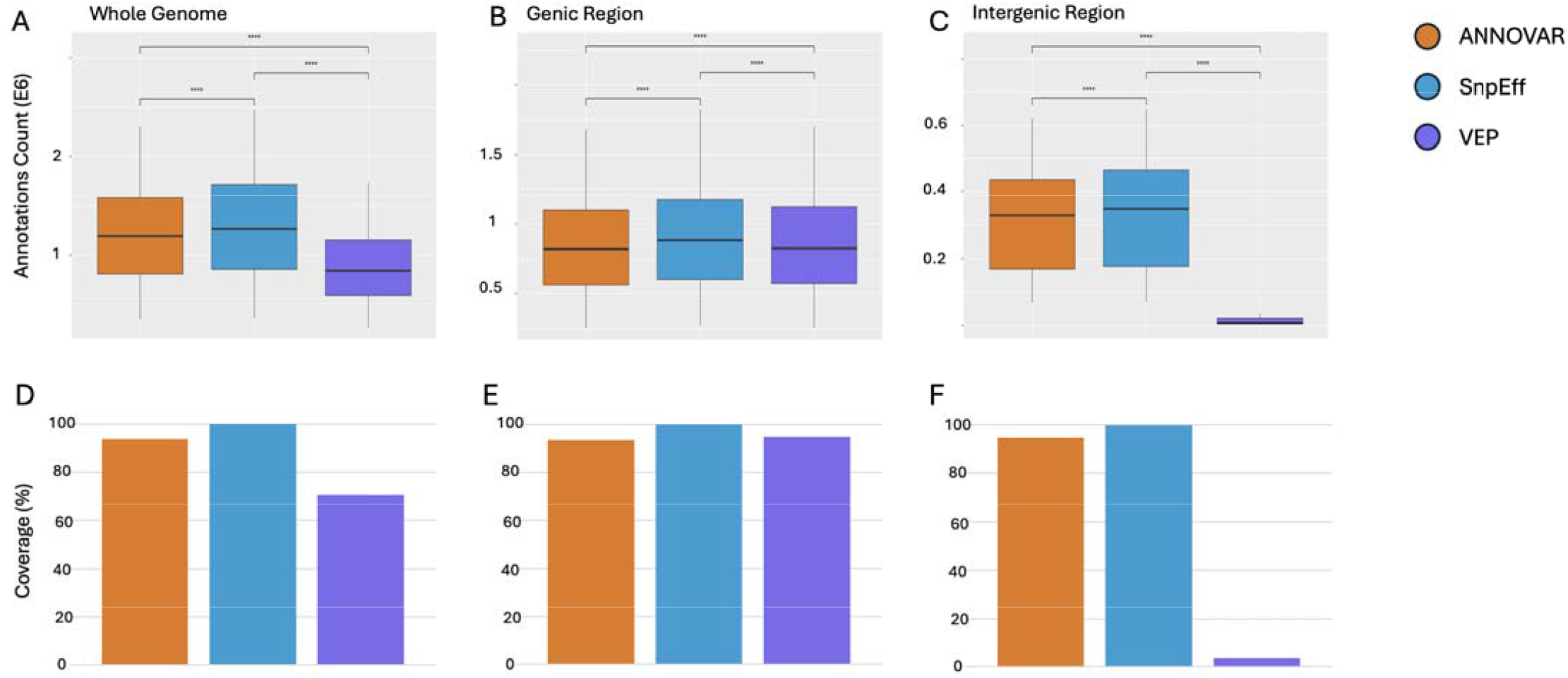
Ensembl Model Comparison. Protein annotation coverage and counts across ANNOVAR, SnpEff, and VEP using the Ensembl gene model. Boxplots (A-C) show the total number of protein annotations per tool across the whole genome (A), genic regions (B), and intergenic regions (C), revealing statistically significant differences across tools (****p-adj < 0.0001). Bar plots (D-F) display the percentage of reference UniProt IDs captured by each tool in the corresponding region, highlighting strong genic performance from all tools and steep intergenic drop-off in VEP.

**Figure 5.**
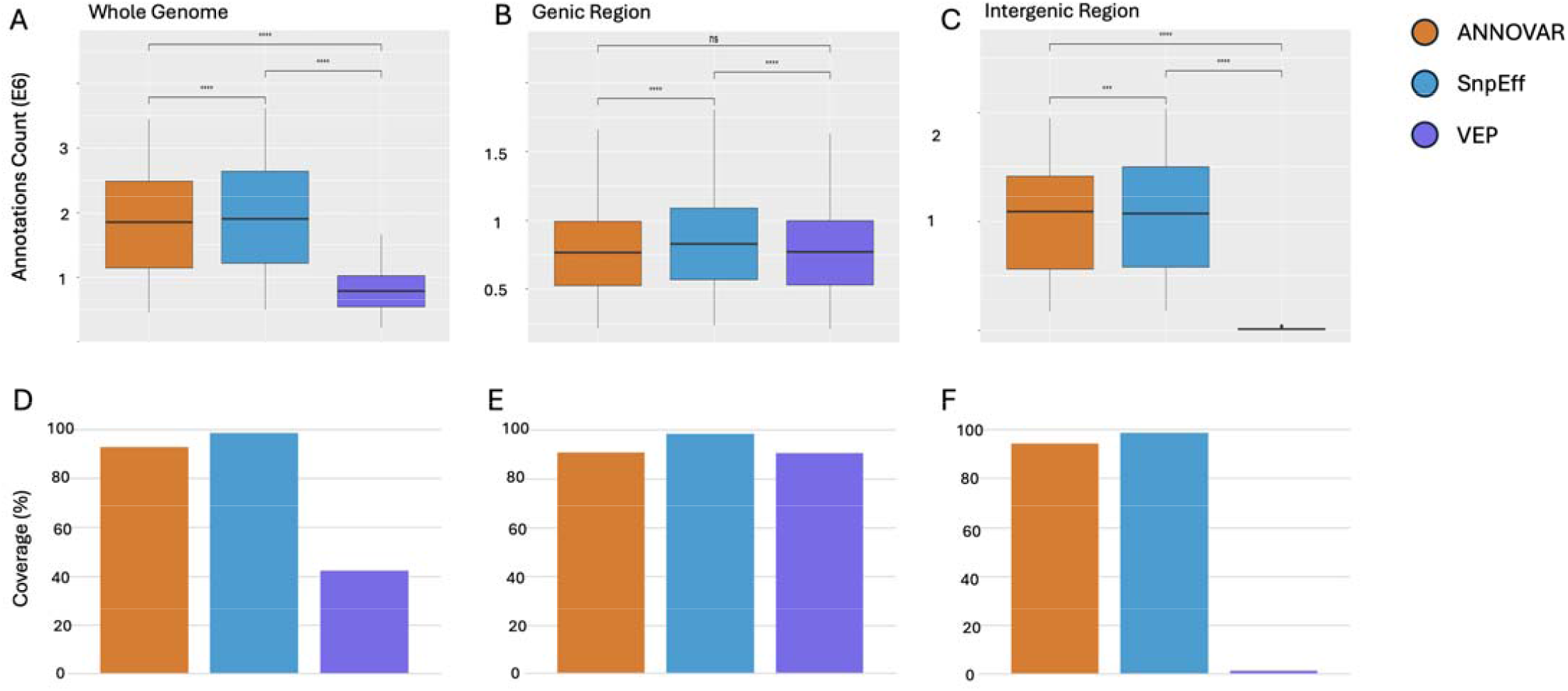
RefSeq Model Comparison. Protein annotation coverage and counts across ANNOVAR, SnpEff, and VEP using the RefSeq gene model. Boxplots (A-C) display the number of protein annotations per tool across the whole genome (A), genic regions (B), and intergenic regions (C). Significant differences were observed across tools in most regions (****p-adj < 0.0001), except between ANNOVAR and VEP in the genic region (ns: p-adj > 0.05; B). Bar plots (D-F) show the proportion of reference UniProt IDs recovered by each tool, with SnpEff achieving near-complete recovery of the union reference and VEP showing major intergenic drop-off.

Within the Ensembl gene model, SnpEff annotated the highest proportion of proteins, capturing 99.9% of the union set genome-wide. ANNOVAR followed closely at 93.6%, while VEP annotated only 70.4% (Fig. 4D). Coverage was high for all tools in genic regions, 100.0% for SnpEff, 93.3% for ANNOVAR, and 94.7% or VEP, but diverged sharply in intergenic regions, where VEP annotated just 3.5% of proteins compared to 99.6% for SnpEff and 94.3% for ANNOVAR (Fig. 4E-F). This divergence is consistent with tool-specific differences in intergenic SNP-to-gene assignment under default configurations.

Using RefSeq, across the genome, the same pattern emerged. SnpEff again achieved the highest coverage (98.5%), followed by ANNOVAR (92.7%) and VEP (42.3%) (Fig. 5D). In genic regions, all three tools had strong performance: 98.5% for SnpEff, 90.8% for ANNOVAR, and 90.6% for VEP (Fig. 5E). However, intergenic coverage again dropped sharply for VEP (1.4%), while SnpEff and ANNOVAR remained above 94% (Fig. 5F).

To compare annotation performance across gene models (Fig. 6; Table 1) while holding the tool constant, we used the standardized region framework defined in the methods. Under this design, each tool’s protein annotations were evaluated using both RefSeq and Ensembl gene models and compared to the combination of both models. Across the whole genome, RefSeq-based annotations yielded higher protein coverage than Ensembl for both ANNOVAR (95.5% vs. 62.9%) and SnpEff (94.8% vs. 62.8%). However, VEP was the sole exception to this pattern, with Ensembl-based annotations achieving higher coverage than RefSeq (97.0% vs. 89.5%). Notably, no single gene model reached 100% coverage, indicating that each contributed unique protein annotations. This trend held across genic and intergenic regions as well, and all differences were statistically significant (p-adj < 0.001), except for the VEP comparison in intergenic regions, which showed no significant difference.

**Figure 6.**
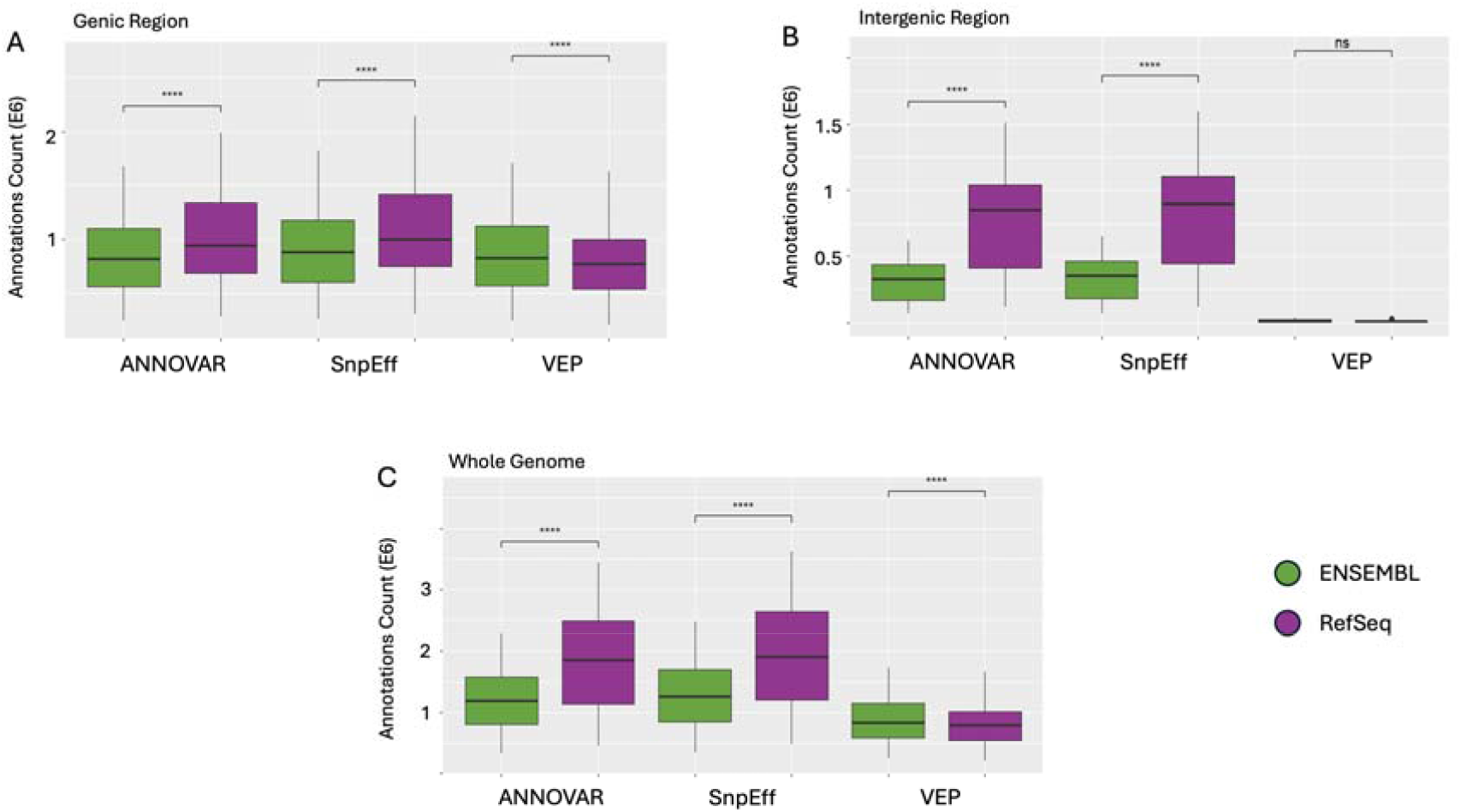
Gene Models Comparison. Protein annotation counts across ANNOVAR, SnpEff, and VEP using Ensembl and RefSeq gene models. (A) Genic region annotations show significantly higher output with RefSeq across all tools (****p-adj < 0.0001). (B) Intergenic annotations are also significantly higher with RefSeq for ANNOVAR and SnpEff, while VEP shows no significant difference (ns). (C) Whole-genome annotations follow the same trend, with RefSeq yielding significantly more annotations than Ensembl across all tools. These results underscore the gene model-dependent variability in protein annotation counts across both genic contexts and tools.

Finally, we evaluated whether integrating tools and gene models created an improvement in protein annotation coverage. Across all regions, combining tools led to significantly more comprehensive protein coverage than using any single tool alone. Combining tools across both Ensembl and RefSeq gene models yielded significantly higher annotation counts than combining tools within a single gene model (p-adj < 0.0001). This pattern held true in genic, intergenic, and whole-genome contexts (Fig. 7)

**Figure 7.**
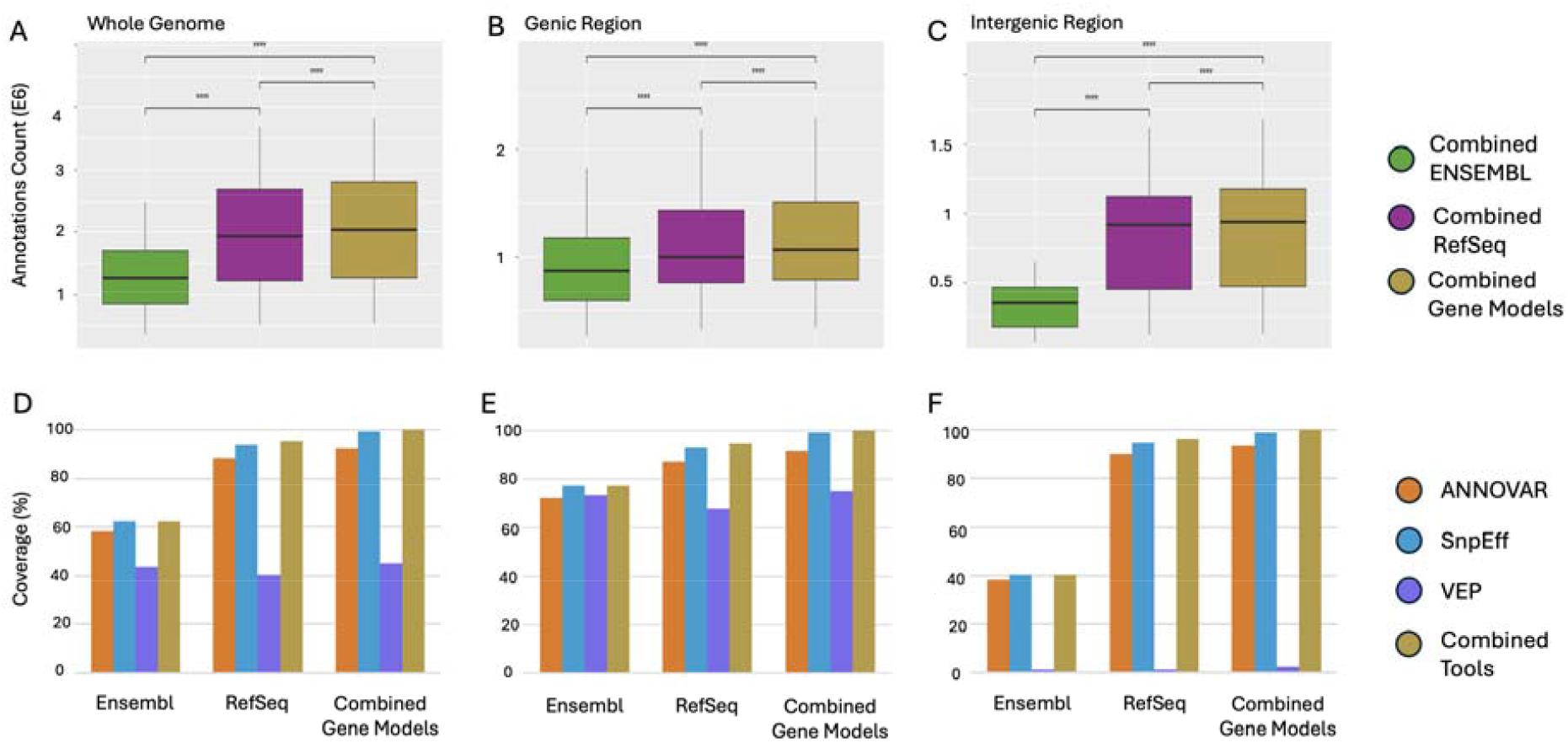
Comparison of annotation performance across different gene models and tool integration strategies. Box plots (A-C) display total annotation counts when using all three tools within Ensembl (“Combined (Ensembl)”), within RefSeq (“Combined (RefSeq)”), or across both models (“Combined Gene Models”). Across the whole genome, genic, and intergenic regions, combining all tools across both models significantly outperforms model-specific combinations in both annotation breadth and coverage (****p-adj < 0.0001). Bar plots (D-F) show the percentage of reference UniProt IDs recovered by each tool and the combined-tool strat gy within RefSeq, Ensembl, and both gene models combined.

Table 2 provides a comprehensive summary of annotation coverage across all combinations of tools and gene models, allowing us to examine how different configurations compare to the fully combined reference set. When comparing RefSeq and Ensembl to the combined gene model output, RefSeq consistently captured substantially more protein annotations for both ANNOVAR and SnpEff, across genic, intergenic, and whole-genome regions. In contrast, VEP performed somewhat better with Ensembl, highlighting that gene model compatibility can vary depending on the tool.

**Table 2.** Protein Annotation Coverage by Tool-Gene Model Combination. Percentage of UniProt IDs captured across all SNPs, stratified by region (whole genome, genic, intergenic) and by annotation tool (ANNOVAR, SnpEff, VEP) within each gene model (RefSeq, Ensembl). Combined tool strategies within each gene model and across both models are also reported. Coverage reflects the proportion of the unified reference set (per region) captured by each configuration. This table highlights substantial variability in annotation depth based on tool and gene model choice, with RefSeq- and multi-tool configurations achieving the highest coverage overall. Accordingly, the fully combined configuration (Combined Tools across Combined gene models) reaches 100% by definition, and the key comparisons are the shortfalls of single-tool and single-gene-model strategies relative to this union reference.

|  | Gene Models |  |  |
| --- | --- | --- | --- |
|  | RefSeq | Ensembl | Combined |
| **Genic** |  |  |  |
| ANNOVAR | 24,138,918 (87.3%) | 20,008,036 (72.4%) | 25,422,674 (92.0%) |
| SnpEff | 25,811,247 (93.4%) | 21,451,709 (77.6%) | 27,461,035 (99.4%) |
| VEP | 18,775,184 (67.9%) | 20,318,587 (73.5%) | 20,777,302 (75.2%) |
| Combined Tools | 26,252,329 (95.0%) | 21,451,709 (77.6%) | 27,636,240 (100.0%) |
| **Intergenic** |  |  |  |
| ANNOVAR | 17,456,051 (90.4%) | 7,394,112 (38.3%) | 18,119,063 (93.9%) |
| SnEff | 18,368,241 (95.1%) | 7,786,423 (40.3%) | 19,128,052 (99.1%) |
| VEP | 226,680 (1.2%) | 275,070 (1.4%) | 451,729 (2.3%) |
| Combined Tools | 18,617,950 (96.4%) | 7,816,748 (40.5%) | 19,305,376 (100.0%) |
| **Whole** |  |  |  |
| ANNOVAR | 41,594,969 (88.6%) | 27,402,148 (58.4%) | 43,541,737 (92.8%) |
| SnEff | 44,179,488 (94.1%) | 29,238,132 (62.3%) | 46,589,087 (99.2%) |
| VEP | 19,001,864 (40.5%) | 20,593,657 (43.9%) | 21,229,031 (45.2%) |
| Combined Tools | 44,870,279 (95.6%) | 29,268,457 (62.3%) | 46,941,616 (100.0%) |

Although SnpEff with the combined gene model achieved 99.2% coverage genome-wide, this configuration did not recover 352,529 union-reference protein annotations, demonstrating that even a single-tool strategy falls short of full coverage.

Importantly, Table 2 shows that only the full integration of all tools and gene models achieved complete recovery of the union reference across all regions (100.0%). This result is expected under the union-based reference definition and highlights the extent to which each single-configuration strategy captures only a subset of the observed annotation breadth. These results suggest that maximal annotation breadth cannot be reached by relying on any single tool or gene model, but rather requires leveraging the diversity of both.

This conclusion is further supported by Figure 7, which compares annotation performance across gene model strategies. Even when tools were combined, configurations based on a single gene model yielded significantly fewer annotations than the fully integrated strategy that combined both tools and gene models (p-adj < 0.0001). This pattern held consistently across all regions, whole genome (Fig. 7A), genic (Fig. 7B), and intergenic (Fig. 7C).

Figures 7D–F present the corresponding coverage percentages for each configuration, based on the values reported in Table 2 for whole genome, genic, and intergenic regions, respectively. These bar plots demonstrate that although combining tools within RefSeq or Ensembl improves coverage, neither gene model alone achieves the completeness provided by integrating all tools and both gene models. For a full statistical comparison of all possible tool and gene model combinations, see Additional File 5.

### Benchmarking Impact on Pathway Interpretation: Case Study Using Colorectal Cancer SNPs

To determine whether these annotation discrepancies influence biological interpretation, we next conducted a pathway enrichment analysis using PANTHER on 204 colorectal cancer–associated SNPs from the FIGI GWAS. We tested different combinations of annotation tools and gene models. When using individual tools with the Ensembl gene model, up to three pathways reached FDR significance: TGF-beta signaling, Alzheimer’s disease–presenilin, and gonadotropin-releasing hormone receptor. In contrast, RefSeq-based annotations enabled the detection of a fourth significant pathway, cadherin signaling, for some tool configurations.

While TGF-beta signaling was consistently enriched across all tools and gene models, results for Alzheimer’s disease–presenilin and cadherin signaling varied. Cadherin signaling was not significant with any tool using Ensembl (FDR ≥ 0.59) and was also missed by VEP when used with RefSeq (FDR = 1.0). Similarly, Alzheimer’s disease–presenilin did not reach significance with VEP and RefSeq (FDR = 0.19). In contrast, both pathways were detected when using RefSeq annotations from ANNOVAR or SnpEff (Additional File 4A).

When all three tools were combined using Ensembl, the same three pathways were detected. Combining the tools with RefSeq identified all four (Additional File 4B). Finally, integrating all three tools and both gene models also captured all four pathways (Additional File 4C).

Although this fully combined strategy did not produce lower FDR values than combining all tools with RefSeq alone, it preserved the complete set of enriched pathways while maximizing annotation coverage. Differences in FDR-adjusted p-values across annotation strategies reflect changes in both the size and composition of the mapped protein sets submitted to PANTHER. In this case study, union-based integration generally produced intermediate significance estimates across pathways, although TGF-beta signaling showed the highest (yet still significant) FDR under full integration. Together, these results suggest that integrating tools and gene models provides a more robust basis for downstream interpretation by prioritizing pathway stability over minimizing FDR values.

### Practical Guidance for Users

For practical use, our results suggest two defensible annotation strategies. Regardless of approach, studies should explicitly report the annotation tool(s) and version(s), the gene model(s) and release(s), and the pathway enrichment resource used (e.g., database name and version) to ensure reproducibility. First, investigators can make an informed single-configuration choice by learning the behavior of their selected annotation tool and gene model, recognizing that these choices can materially change downstream protein sets and enrichment results. Alternatively, when the goal is to maximize robustness to tool- and gene model– specific variability, a conservative default is to integrate across configurations by taking the union of annotations across tools and/or gene models, as we show this approach recovers the broadest set of gene and protein annotations and stabilizes downstream pathway inference. This integration can be implemented either locally, if users already maintain multiple annotators, or without local installation by using AnnoQ to query multiple tool–gene model outputs and then applying the union logic. To support reproducibility and adoption, we provide scripts that demonstrate the exact logic used to extract and standardize gene identifiers from SNP-level outputs and map them through HGNC-linked identifiers for consistent downstream comparisons and enrichment.

## DISCUSSION

This study reveals substantial variability in SNP-to-protein annotations driven by both the annotation tools and the reference gene model used. No single tool–gene model combination achieved complete recovery of the union reference, and our genome-wide evaluation underscores how these discrepancies propagate through to the protein level. By systematically applying both a qualitative analysis of annotation overlap patterns and a quantitative analysis of annotation completeness, we highlight not only where tools diverge but also how those differences influence downstream biological interpretation.

One of the most striking findings was the large discrepancy in protein annotation output between RefSeq and Ensembl: RefSeq mapped ∼32.6% more SNPs to at least one protein than Ensembl (34,349,476 vs 25,905,898) and produced ∼53.3% more SNP-to-protein annotation events (44,870,279 vs 29,268,457; Figure 1). This pattern held across both genic and intergenic regions and remained consistent even after region definitions were standardized to those of Ensembl. While Ensembl exhibited greater internal agreement among tools, RefSeq consistently annotated more proteins, particularly in intergenic regions, suggesting a broader, more permissive annotation scope (Table 1).

Annotation tool coverage patterns varied depending on genomic context. SnpEff demonstrated high and consistent coverage across all configurations, making it the most robust performer in our comparison. VEP, in contrast, performed well in genic regions but showed steep drop-offs in intergenic coverage, particularly when used with RefSeq (Figures 4 - 6). ANNOVAR showed intermediate performance across most conditions.

The practical impact of these tools and gene model differences is substantial. As shown in Table 2, genome-wide protein coverage ranged from as low as 40.5% (VEP with RefSeq) to as high as 94.1% (SnpEff with RefSeq). Even top-performing combinations differed by several hundred thousand annotations. While using SnpEff across both gene models achieved 99.2% coverage, this still left over 352,529 proteins uncaptured, reinforcing the additive value of full integration. When tools were combined within a single gene model, coverage improved (Figure 7A-C), but combining tools across gene models consistently yielded significantly greater coverage than any within-model configuration (Figure 7D-F, Table 2).

These annotation differences had meaningful downstream consequences in our colorectal cancer case study. When using individual tools with the Ensembl gene model, up to three pathways reached statistical significance: TGF-beta signaling, Alzheimer’s disease–presenilin, and gonadotropin-releasing hormone receptor. RefSeq-based annotations enabled the detection of a fourth pathway, cadherin signaling, depending on the tool used. However, both cadherin signaling and Alzheimer’s disease–presenilin were missed when using VEP with RefSeq (FDR = 1.0 and 0.19, respectively), highlighting the variability introduced by annotation choices. In contrast, both pathways were successfully detected when using RefSeq annotations from ANNOVAR or SnpEff.

Combining all three tools with Ensembl recovered the same three pathways, while combining tools with RefSeq yielded all four. The fully integrated approach, merging all tools and both gene models, also detected all four pathways. Although this strategy did not improve statistical significance beyond combining all tools with RefSeq and in some cases produced slightly higher but still significant FDR values, it ensured the broadest annotation coverage and preserved all enriched pathways. These results underscore the importance of using both multiple tools and multiple gene models to support more comprehensive and reliable downstream interpretation.

Importantly, differences in pathway significance across annotation strategies do not reflect uncertainty in the pathway enrichment test itself, which is well defined statistically, but rather uncertainty in how SNPs are mapped to genes prior to enrichment analysis. Because there is no gold standard for selecting or prioritizing annotation tools or gene models, different strategies yield distinct but internally valid gene sets that define different enrichment hypotheses. Our results suggest that integrating annotations across tools and gene models, or explicitly evaluating pathway stability across strategies, provides a more robust framework for interpretation than relying on any single annotation configuration.

In practice, we suggest using an integrated, combined annotation strategy as a default, or explicitly evaluating pathway stability across tools and gene models when a single configuration is used.

## CONCLUSION

This genome-wide assessment of SNP annotation consistency demonstrates that both tool and gene model selection substantially influence protein-level outputs. RefSeq produced more total annotations, particularly in intergenic regions, while Ensembl showed stronger agreement across tools. SnpEff consistently delivered high coverage across all contexts, whereas VEP’s performance was more variable, especially in intergenic regions. Coverage varied widely across tool–gene model combinations, with Table 2 illustrating how these choices can shift annotation outcomes by millions of SNP-to-protein annotations.

These findings highlight the importance of deliberate annotation strategies in genomic research. Our colorectal cancer pathway case study showed that upstream choices in tools and gene models can directly affect downstream biological interpretation. While no single combination was sufficient, the fully integrated approach, merging all tools and both gene models, captured the broadest set of annotations and recovered all significant biological pathways. Although this strategy may introduce larger gene sets and modestly attenuate statistical significance, its advantages were consistent across genic, intergenic, and whole-genome analyses. Together, these results advocate for a multi-tool, multi-gene model strategy as a robust default for pathway analysis, with genomic context guiding interpretation rather than tool or model selection.

Although integrating multiple tools and gene models can increase analytical complexity, this represents a practical rather than conceptual limitation. Future work will focus on streamlining multi-tool, multi-model annotation strategies to facilitate broader adoption.

## METHODS

Because SNP annotation can yield multiple gene and protein IDs per variant, we first describe how outputs were standardized and then how agreement was evaluated.

### Data Sources and Annotation Tools

This study was conducted on the AnnoQ platform^29^ (version 1.11) and used single-nucleotide polymorphism (SNP) data from the Haplotype Reference Consortium (HRC.r1-1) dataset; annotations were generated within the AnnoQ environment (WGSA v0.95)^30^. All analyses were performed on hg38/GRCh38 coordinates with HRC variants processed in hg38 coordinates within the AnnoQ/WGSA workflow. Gene-model annotations were generated using Ensembl release 107 (consistent with GENCODE v41) and the RefSeq gene model bundled within the AnnoQ/WGSA v0.95 environment. Annotations were retrieved from three commonly used tools, ANNOVAR (2020-06-07), SnpEff version 4.3t, and VEP version 107, through the AnnoQ interface (https://annoq.org). AnnoQ executes each tool with its standard, tool-recommended configuration for the specified gene model; we did not impose cross-tool harmonization of gene boundary definitions, upstream or downstream distance thresholds, or intergenic assignment rules. As a result, differences observed across tools reflect tool-native behavior rather than post hoc parameter tuning. PANTHER 19.0 was used for pathway enrichment analyses. All analyses were conducted using custom Python^31^ (version 3.12) and R^32^ (version 4.4.1) scripts and data available at https://github.com/USCbiostats/SNP-Annotation-Agreement-and-Downstream-Impact-Analysis.

### Region Classification: Genic and Intergenic SNPs

SNPs were categorized as genic or intergenic using ANNOVAR functional annotation outputs under Ensembl and RefSeq gene models. This stratification was used only for context-specific reporting and statistical aggregation; it does not alter or constrain tool outputs, and it should not be interpreted as a tool-provided ground-truth region label. Genic regions were defined as overlapping exons, introns, or untranslated regions (UTRs), while intergenic SNPs were those located outside annotated gene boundaries and linked to the nearest gene. ANNOVAR was selected for this step because its output clearly distinguishes between the two categories: genic annotations are listed under the “ANNOVAR Gene ID” field, and intergenic ones under the “ANNOVAR Closest Gene ID” field. Because ‘genic’ and ‘intergenic’ depend on gene model boundaries, we classified SNPs using each gene model’s boundaries for within-model tool comparisons. When directly comparing gene models, we used fixed boundaries (Ensembl) to keep region definitions constant across comparisons and to avoid combining gene model boundary differences with tool-specific annotation behavior in the stratified analyses.

### ID Mapping

To enable consistent protein-level comparisons across tools and gene models, we mapped gene-level annotations to UniProt^33^ identifiers. Depending on the gene model, we used gene symbols and Entrez IDs^34^ for RefSeq-based annotations, and Ensembl gene IDs for Ensembl-based annotations, consistent with how each gene model defines its identifiers.

Because PANTHER enrichment operates on identifiers that map to PANTHER protein records, we restricted analyses to protein-coding genes. Pseudogenes and non-protein-coding loci were excluded because they do not consistently map to PANTHER protein entries and therefore are not reliably testable in this enrichment framework.

To standardize identifiers across tools and gene models, we used the HUGO Gene Nomenclature Committee (HGNC)^35^ approved gene dataset^36^, which links gene symbols, Entrez IDs, and Ensembl gene IDs to UniProt IDs. This file was used to enforce a single gene-to-UniProt mapping (one UniProt ID per gene) to avoid inflation from transcript isoforms. Applying this standardized mapping step ensured comparability across annotations and supported downstream pathway analysis.

To verify that the mapping process did not introduce discrepancies, we applied it uniformly across all tools and gene models. During this process, we identified a number of outdated Ensembl gene IDs missing from the HGNC file. To confirm these were deprecated and not mistakenly excluded, we manually checked a subset in Ensembl and verified that the missing IDs had been superseded.

### Annotation Agreement Framework

Because no gold standard exists for SNP-to-protein annotation, we treated the union of all observed annotations as a pragmatic reference.

Agreement across tools and gene models was evaluated using a SNP-level comparison framework at the protein level. All comparisons were based on UniProt IDs derived from the standardized gene annotations described in the previous ID Mapping section. For each SNP, we constructed a reference set of UniProt IDs by taking the union of all protein annotations returned by the tools being compared. This reference set served as the basis for both qualitative and quantitative analyses. For example, if ANNOVAR returns proteins {A, B}, SnpEff returns {A, C}, and VEP returns {A, B, C} for the same SNP, the reference set is {A, B, C}. A tool is considered in full agreement only if it matches {A, B, C}, and in partial agreement if it matches a subset.

First, we conducted a **qualitative analysis** to characterize patterns of annotation agreement. For each SNP, each tool or gene model was assessed relative to the reference set and classified as exhibiting full agreement if it recovered the complete reference set, partial agreement if it recovered a subset of reference proteins, or no agreement if it recovered none. SNPs were then assigned to categorical classes based on the combination of agreement outcomes observed across tools or gene models.

At the tool level, categories included full agreement across all three tools, tool-specific matches (e.g., only SnpEff fully recovering the reference set), pairwise agreement cases in which two tools fully recovered the reference set while the third did not, nested subset cases in which two tools fully recovered the reference set while the third recovered only a subset, and fully disjoint cases in which no tool fully recovered the reference set. In total, 17 distinct tool-level agreement categories were observed (Additional File 2).

At the gene model level, SNPs were classified into seven categories by comparing RefSeq- and Ensembl-derived UniProt identifiers relative to the SNP-specific reference set. For each SNP, each gene model was evaluated according to whether it fully recovered the reference set, recovered only a subset of reference proteins, or contributed no unique annotations. Based on these outcomes, SNPs were assigned to one of seven categories, including Exact Match (both gene models fully recovering the reference set), RefSeq with Ensembl Subset, Ensembl with RefSeq Subset, partial overlap between gene models, fully disjoint annotations, and gene-model-specific cases in which only RefSeq or only Ensembl contributed annotations (Additional File 3).

Second, we performed a **quantitative analysis** to assess annotation completeness per SNP. For each tool, we recorded the number of UniProt IDs it captured out of the total in the SNP’s reference set. These numerator-denominator pairs were retained for two distinct purposes: the numerators were used to perform pairwise statistical comparisons across tools and gene models, while the denominators were used to calculate coverage percentages (Tables 1 & 2).

In a complementary analysis, we also evaluated how annotation output varied when holding the tool constant and switching gene models. This comparison allowed us to directly measure the effect of gene model selection independently from tool behavior and served as an intermediate step between tool-based and fully integrated analyses.

The qualitative analysis reveals which tools disagree, while the quantitative analysis measures how much annotation is lost.

A schematic overview of this framework is provided in Figure 3.

### Statistical Analysis

To compare total protein annotation output across tools, we aggregated annotation counts by chromosome. For each chromosome (1–22 and X), we recorded the number of protein-level annotations produced by each tool based on SNPs located on that chromosome.

Although annotations were evaluated at the SNP level to characterize agreement patterns, formal statistical testing was performed on chromosome-level aggregates rather than individual SNPs to avoid SNP-level hypothesis testing that would incorrectly treat linkage disequilibrium-correlated variants as independent samples and to limit inflation of statistical significance.

We used paired t-tests to compare annotation counts between tools, treating chromosomes as the matched units. Bonferroni correction was applied for pairwise comparisons (ANNOVAR vs. SnpEff, ANNOVAR vs. VEP, and SnpEff vs. VEP). These tests were performed separately for genome-wide, genic, and intergenic regions. For comparisons between gene models (RefSeq vs. Ensembl), Bonferroni correction was also applied. Box plots and bar plots were used to visualize annotation counts and tool comparisons (Figures 4-7).

Proportion-based tests were not used because the true number of expected protein annotations per SNP is unknown. Chromosomes were treated as matched units (n = 23) because annotation pipelines were applied uniformly across all chromosomes, enabling consistent aggregation and comparison of annotation outputs across tools, gene models, and their combinations.

Counts represent SNP-to-protein annotation events (i.e., a SNP annotated to multiple proteins contributes multiple annotations).

### Impact on Pathway Interpretation: Case Study Using Colorectal Cancer SNPs

To illustrate how annotation tools and gene model selection can affect downstream interpretation, we conducted a case study using 204 SNPs from a genome-wide association study (GWAS) of colorectal cancer conducted by the Functionally Informed Gene–Environment Interaction (FIGI) consortium^27^. Each SNP was annotated using ANNOVAR, SnpEff, and VEP, with both Ensembl and RefSeq gene models, generating six single-tool configurations (3 tools × 2 gene models; Additional File 4A). We then evaluated two integrative strategies: (i) within-gene-model integration, in which UniProt IDs were defined as the union of unique mapped proteins across all three tools within Ensembl or within RefSeq (Additional File 4B), and (ii) full integration, in which UniProt IDs were defined as the union of unique mapped proteins across all tools and both gene models (Additional File 4C). No prioritization rule was applied when annotations disagreed; combined strategies retained all unique mapped proteins.

For each configuration, we performed pathway enrichment analysis using the PANTHER 19.0 Classification System^11^ (pantherdb.org), with False Discovery Rate (FDR) correction for multiple testing across pathways, to compare how the annotation strategy altered the set of significant pathways^28^. We used PANTHER’s overrepresentation test (Fisher’s exact test) with the *Homo sapiens* reference list. The input list consisted of UniProt IDs derived from protein-coding genes, consistent with the annotation agreement framework.

## DATA AVAILABILITY

All analyses were performed using the publicly available data aggregated in the AnnoQ platform (annoq.org)^29^. Single-nucleotide polymorphism (SNP) data is from the Haplotype Reference Consortium (HRC.r1-1) dataset (https://ega-archive.org/datasets/EGAD00001002729). Gene-model annotations were generated using Ensembl release 107 (https://jul2022.archive.ensembl.org/index.html) and the RefSeq gene model (https://ftp.ncbi.nlm.nih.gov/refseq/H_sapiens/) bundled within the AnnoQ. No new sequence or genetic variation data were generated in the analysis. The scripts for the analysis are available in a GitHub repository described in the Methods section of the manuscript.

## ACKNOWLEDGEMENTS

The authors are grateful to Paul Majoram and Michele Ramos Correa for helpful discussions and comments to the manuscript.

## FUNDING

This work was supported by the National Cancer Institute (NCI) grant P01-CA196569.

**Ethics, Consent to Participate, and Consent to Publish declarations**: not applicable

